# CINmetrics: An R package for chromosomal instability analysis

**DOI:** 10.1101/2021.11.15.467294

**Authors:** Vishal H. Oza, Jennifer L. Fisher, Roshan Darji, Brittany N. Lasseigne

## Abstract

Genomic instability is an important hallmark of cancer and more recently has been identified in others like neurodegenrative diseases. Chromosomal instability, as a measure of genomic instability, has been used to characterize clinical and biological phenotypes associated with these diseases by measuring structural and numerical chromosomal alterations. There have been multiple chromosomal instability scores developed across many studies in the literature; however, these scores have not been compared because of the lack of a single tool available to calculate and facilitate these various metrics. Here, we provide an R package CINmetrics, that calculates six different chromosomal instability scores and allows direct comparison between them. We also demonstrate how these scores differ by applying CINmetrics to breast cancer data from The Cancer Genome Atlas (TCGA). The package is available on CRAN at *https://cran.r-project.org/package=CINmetrics* and on github at *https://github.com/lasseignelab/CINmetrics*.

## INTRODUCTION

Genomic instability, one of the hallmarks of cancer and aging, is measured in many forms such as chromosomal instability, microsatellite instability, and instability characterized by increased frequency of base-pair mutations (Bakhoum and Cantley, 2018; Pikor et al., 2013; Negrini et al., 2010; López-Otín et al., 2013). Particularly, chromosomal instability (CIN) is associated with cancer progression, tumor immunity, and inflammation (Pikor et al., 2013; Bach et al., 2019). Recently, CIN has been shown to contribute to diseases other than cancer, including neurodegenerative diseases (Hou et al., 2017; Yurov et al., 2019).

CIN is broadly defined as the change in number and structure of chromosomes (Vargas-Rondón et al., 2017). CIN has been measured directly from tumor specimens by capturing errors during anaphase segregation or indirectly by measuring numerical and structural chromosomal alterations across cell populations. There have been various CIN scores developed across multiple studies involving different cancers which calculate numerical and structural alterations in the chromosome (McGranahan et al., 2012). The differences in calculation of these scores have been associated with different clinical and biological phenotypes (Baumbusch et al., 2013; Davison et al., 2014; Roylance et al., 2011; Bonnet et al., 2012); however, there has been no systematic comparison of different CIN scores and how they vary across and within different cancers. The primary reason being lack of availability of computational framework to calculate these CIN scores. While other packages are available to calculate chromosomal instability (Song et al., 2017), they are limited to a single chromosomal instability score and do not provide a framework to calculate and compare other CIN scores. Here, we provide an R package that provides a unified framework to calculate multiple CIN metrics on same dataset. This package will accelerate chromosomal instability studies by facilitating score comparisons across cancers or other diseases.

## METHODS

The chromosomal instability metrics were mined from the cancer literature and implemented as functions in our CINmetrics R package, based on their ability to detect either structural, numerical, or whole genome instability. The six functions (*tai, taiModified, cna, countingBreakPoints, countingBaseSegments, fga*) are outlined below based on the similarity of the algorithms used to calculate them.

### Total aberration index *(tai)* and Modified Total Aberration Index *(taiModified)*

Total Aberration Index (TAI) was proposed by (Baumbusch et al., 2013) to measure the genomic aberrations in serous ovarian cancers. TAI calculates absolute area under the curve for a copy number segment profile generated by piecewise constant fitting (PCF) algorithm (Baumbusch et al., 2008). Biologically, TAI can be interpreted as absolute deviation from the normal copy number state averaged over all genomic locations. TAI provides a numerical measure in terms of both prevalence as well as the genomic size of copy number variations in tumors. One of the limitations of TAI is that since it was designed for studying advanced stage ovarian tumors, short aberrations found in early stage tumors have low impact on TAI. Therefore, TAI should be used to study the global scale genomic disorganization most likely to occur in late stage tumors.

*tai* implemented in CINmetrics takes into account only those sample values that are in aberrant copy number state, i.e., has a mean segment value of less than or equal to −0.2 and greater than or equal to +0.2.

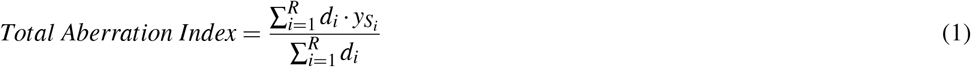

where 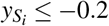 and 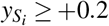 represents the mean segment value, *d*_*i*_ represents the segment length, and *R* represents the total number of segments. Alternatively, *taiModfied* takes into account all the mean segment values and thus preserves the “directionality” of the score.

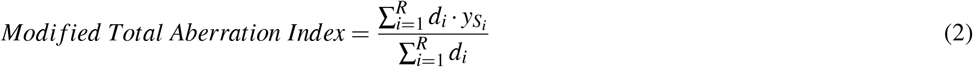

where 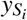 represents the mean segment value, *d*_*i*_ represents the segment length, and *R* represents the total number of segments.

### Copy number abnormality *(cna)* and number of break points *(countingBreakPoints)*

Copy number abnormality (CNA) was developed by (Davison et al., 2014) for studying aneuploidy in superficial gastroesophageal adenocarcinoma. An individual CNA is defined as the segment with copy number outside the predefined range of 1.7 to 2.3 where 2 indicates no loss or gain (assuming that the tumor is diploid) as determined by the Partek segmentation algorithm (Grayson and Aune, 2011). Total CNA for the sample can thus be defined as total number of individual CNAs. CNA represents a measure of segmental aneuploidy. *cna* implemented in CINmetrics is similar except we define individual CNA as the segment with copy number less than or equal to −0.2 and greater than or equal to +0.2 with segment mean of 0 indicating no loss or gain. We chose ± 0.2 as a conservative cutoff for TCGA data as described in (Laddha et al., 2014). The users can modify the cutoff by modifying *segmentMean* parameter.

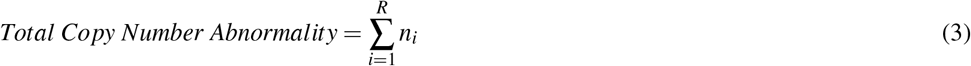

where *n*_*i*_ represents number of segments with 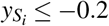 and 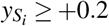, *R* represents the total number of segments and the minimum segment length *d*_*i*_ is greater than or equal to 10.

*countingBreakPoints* is similar to the total breakpoints implemented in (Lee et al., 2011). Segments with mean less than or equal to − 0.2 and greater than or equal to +0.2 and that contain a number of probes above the user defined threshold, are counted and then the value is doubled to account for 3’ and 5’ ends. This metric yields similar results to *cna*

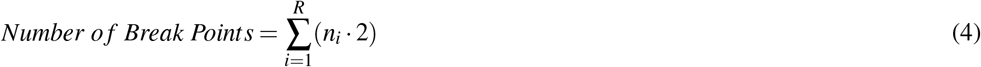

where *n*_*i*_ represents number of segments with 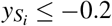 and 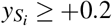 and *R* represents the total number of segments.

### Counting altered base segments *(countingBaseSegments)* and the fraction of the genome altered *(fga)*

Counting altered base segments and fraction of the genome altered are modified implementations of the Genome Instability Index (GII) as described in (Chin et al., 2007). The GII was computed in two different ways, both based on calculating common regions of alteration (CRA). These approaches show high concordance.

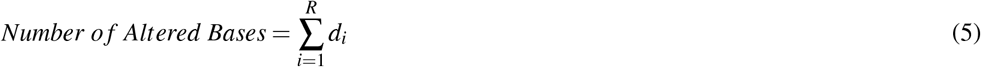

where *d*_*i*_ represents length of segments with 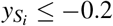 and 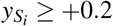 and *R* represents the total number of segments.

*fga* implemented in our package is based on identifying CRAs as fraction of the genome altered. Therefore, the *fga* values are normalized by dividing it by the length of the genome covered. *countingBaseSegments* on the other hand calculates the CRAs.

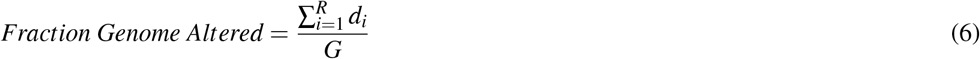

where *d*_*i*_ represents length of segments with 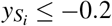 and 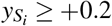, *G* represents genome length covered, and *R* represents the total number of segments. The default value is calculated based on length of each probe on Affeymetrix 6.0 array and excludes the sex chromosomes. One important difference to note is that in the original GII calculations, the algorithm merges the non-overlapping regions between samples, whereas *fga* and *countingBaseSegements* implemented in CINmetrics package do not merge non-overlapping regions.

## RESULTS

We used harmonized masked copy number segment data for breast cancer (BRCA) from The Cancer Genome Atlas (Cancer Genome Atlas Network, 2012; Cerami et al., 2012) to visualize and compare the chromosomal instability metrics implemented in the CINmetrics package. We chose the breast cancer data as it has been shown to exhibit chromosomal instability and thus provides a robust dataset for applying CINmetrics e.g. (Duijf et al., 2019; Voutsadakis, 2021). Figure 1A shows the distribution of CINmetrics in BRCA data for normal and tumor samples. The metrics have been log_10_ scaled to allow for comparison between them. *cna, countingBreakPoints, fga*, and *countingBaseSegments* show an overall pattern of increased genomic instability in tumor samples compared to normal. However, the difference in mean and standard deviation between the two classes (normal and tumor) is very different between these metrics. *tai* and *taiModified* do not capture this global pattern of difference between normal and tumor samples. As mentioned earlier, *tai* and *taiModified* are best suited for late stage cancers, thus should be used as a measure for studying overall genomic disorganization in individual patients with advanced tumors and not as a measure of genomic instability comparison between normal and tumor samples, as we further demonstrate here.

**Figure 1.**
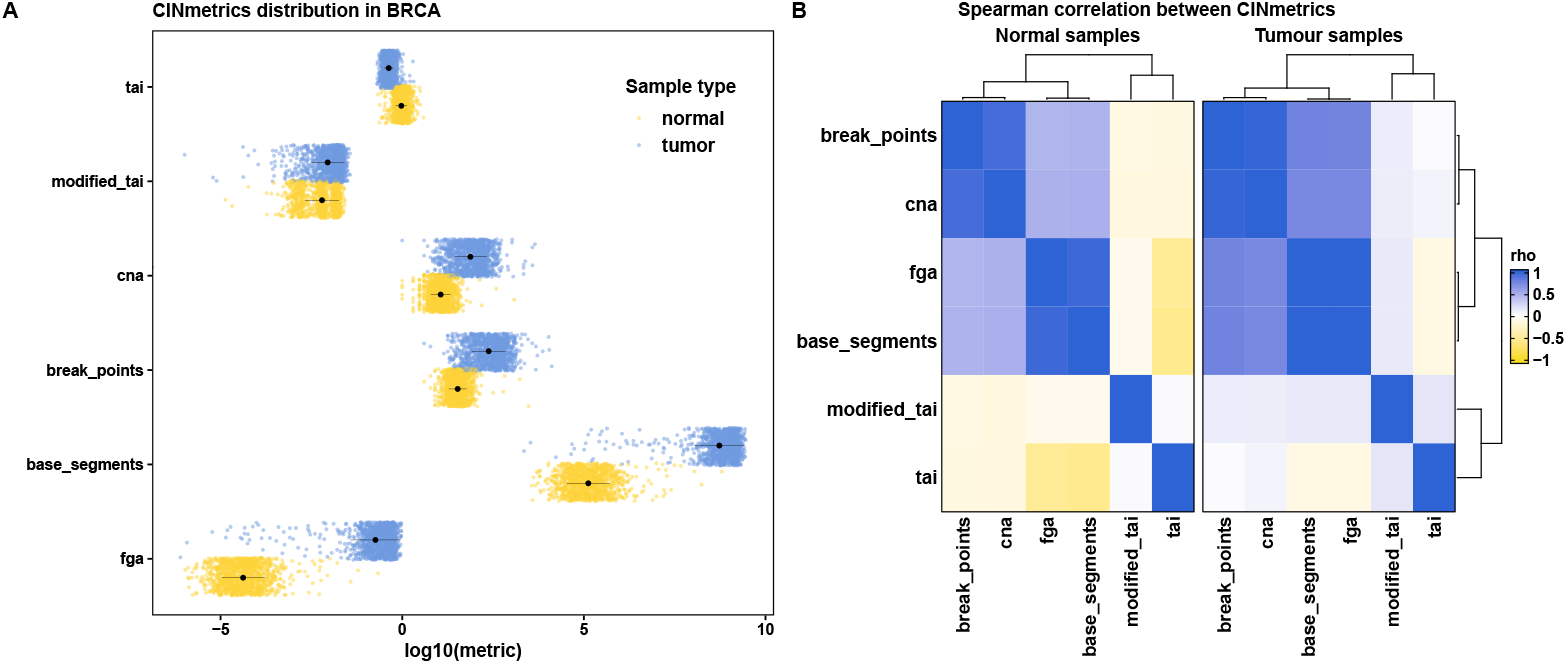
CINmetrics applied to the BRCA dataset from TCGA, A) the distribution of metrics between normal (yellow) and tumor (blue) samples, where the black dot indicates the mean and the black line indicates the standard deviation and B) heatmap of the spearman correlation and complete linkage clustering of the metrics in normal and tumor samples.

To further understand and characterize the relationship between various metrics implemented in CINmetrics, we performed spearman correlation (Spearman, 1904), followed by complete linkage clustering (Vijaya et al., 2019) as shown in Figure 1B. This clustering further demonstrated that *cna, countingBreakPoints, fga*, and *countingBaseSegments* are more similar and therefore highly correlated compared to *tai* and *taiModified*. Furthermore, this relationship is preserved in both normal and tumor samples indicating the four metrics show consistent results and can be used for comparing genomic instability between the two conditions.

We also looked at how the metrics are affected by potential confounders such as tumor purity and ploidy levels in tumor samples in the BRCA dataset. We obtained the purity and ploidy data for BRCA calculated using ABSOLUTE algorithm (Carter et al., 2012) from the NIH Genomic Data Commons Portal (The Pan-Cancer Atlas, 2022).

For purity, the purity score had a range between 0 and 1 for each sample, with 1 being the highest. We divided the score in four quantiles and plotted the density of the samples in each quantile against the CINmetrics scores (Figure 2). All CINmetrics had relatively lower scores for samples with less purity (1st Quantile). Interestingly, *tai* showed a distinct increase in the score with the increase in purity of samples, thus higher purity samples had higher chromosomal instability. *taiModified* showed the same distribution across the four quantiles.

**Figure 2.**
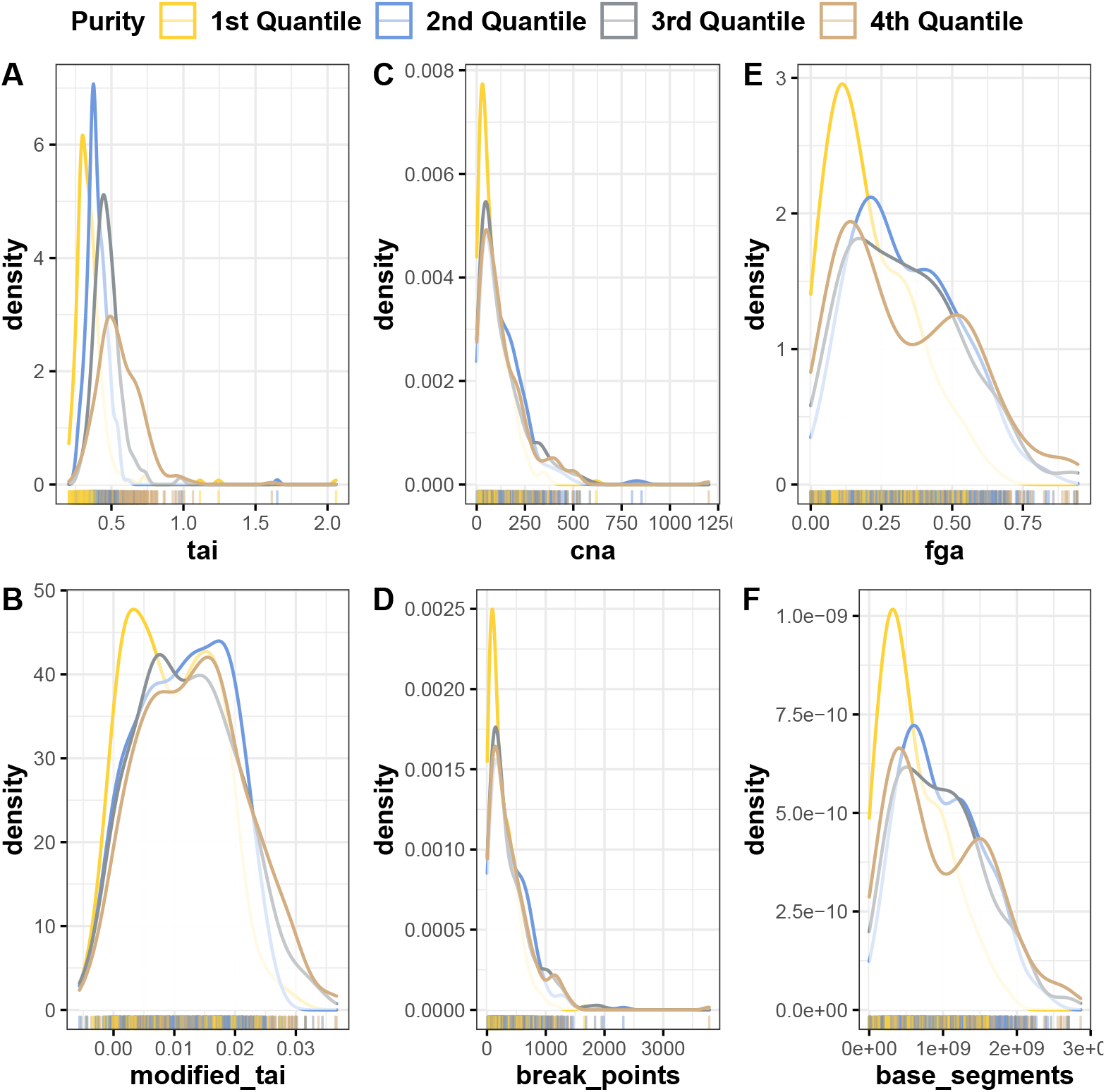
Distribution of CINmetrics applied to BRCA dataset from TCGA compared to sample purity, A) *tai*, B) *taiModified*, C) *cna*, D) *countingBreakPoints*, E) *fga*, and F) *countingBaseSegments*.

For ploidy, we looked at the density of samples with different ploidy numbers against the CINmetrics scores (Figure 3). Samples that were diploid (2n) had the lowest score across *cna, countingBreakPoints, fga*, and *countingBaseSegments*; however not in *tai* and *taiModified. cna* and *countingBreakPoints* were developed to study aneuploidy (Davison et al., 2014; Lee et al., 2011), and they show higher scores corresponding to higher ploidy levels.

**Figure 3.**
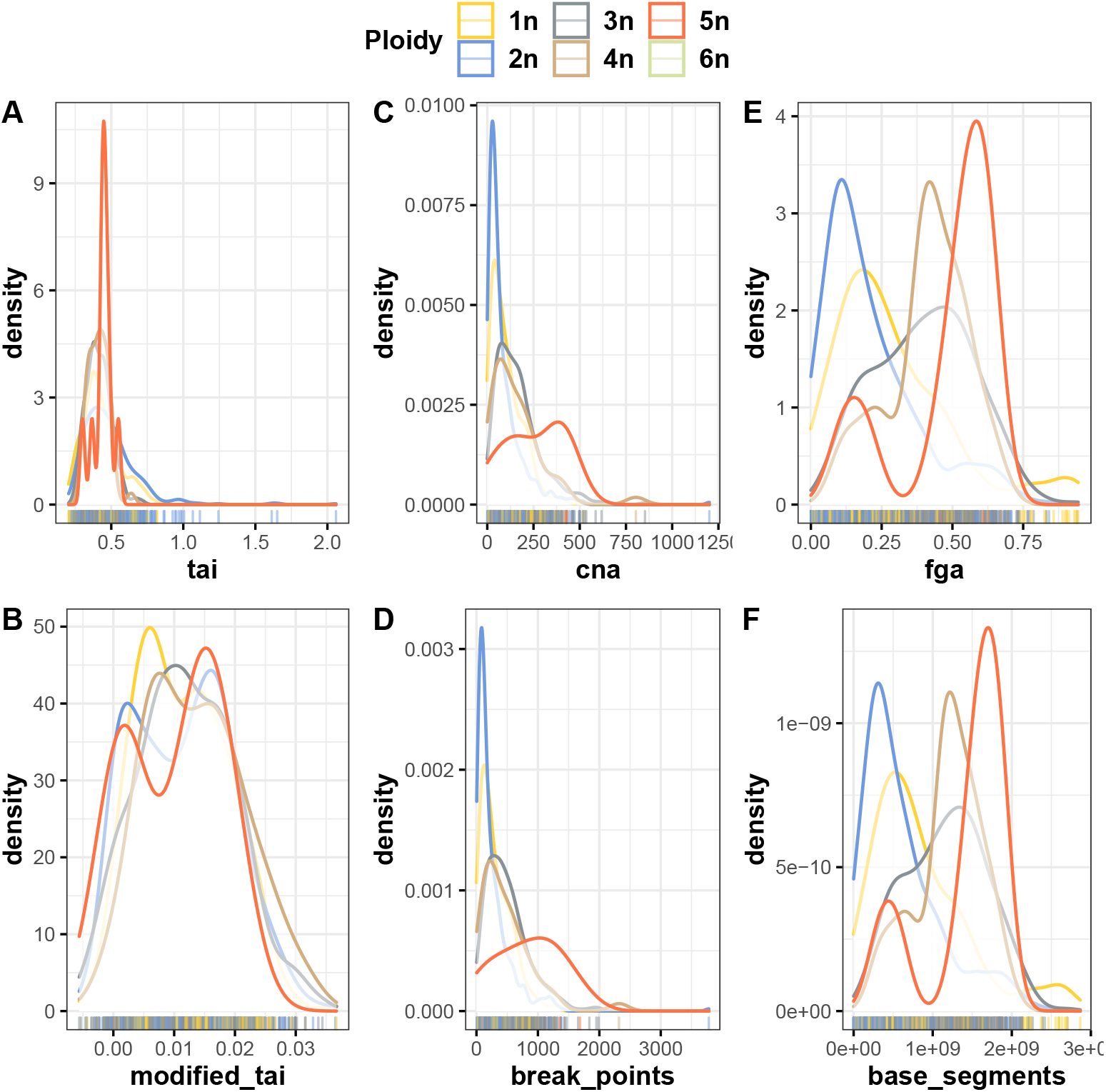
Distribution of CINmetrics applied to the BRCA dataset from TCGA compared to sample ploidy levels, A) *tai*, B) *taiModified*, C) *cna*, D) *countingBreakPoints*, E) *fga*, and F) *countingBaseSegments*.

## CONCLUSIONS

CIN has been one of the most important factors in understanding disease etiology and progression in cancer (Pikor et al., 2013; Bach et al., 2019) and is becoming increasingly recognized for others like neurodegenerative diseases (Hou et al., 2017; Yurov et al., 2019). Numerous methods have been developed to quantitate and characterize the role of chromosomal instability in specific cancers, however, lack of comprehensive tools that calculates these metrics has limited direct comparison between them. Here, we have collected chromosomal instability metrics from the literature and provide them as an R package and associated vignette that allows for reproducible calculations and comparisons. Further, we used BRCA data from The Cancer Genome Atlas to show how the metrics relate to each other. This package thus, provides a useful framework to better characterize and understand genomic instability in cancer.

## ACKNOWLEDGMENTS

We would like to thank Tabea M. Soelter and other members of the Lasseigne Lab for their valuable input.

## REFERENCES

Bach, D.-H., Zhang, W., and Sood, A. K. (2019). Chromosomal instability in tumor initiation and development. Cancer Res., 79(16):3995–4002.

Bakhoum, S. F. and Cantley, L. C. (2018). The multifaceted role of chromosomal instability in cancer and its microenvironment. Cell, 174(6):1347–1360.

Baumbusch, L. O., Aarøe, J., Johansen, F. E., Hicks, J., Sun, H., Bruhn, L., Gunderson, K., Naume, B., Kristensen, V. N., Liestøl, K., Børresen-Dale, A.-L., and Lingjaerde, O. C. (2008). Comparison of the agilent, ROMA/NimbleGen and illumina platforms for classification of copy number alterations in human breast tumors. BMC Genomics, 9:379.

Baumbusch, L. O., Helland, Å., Wang, Y., Liestøl, K., Schaner, M. E., Holm, R., Etemadmoghadam, D., Alsop, K., Brown, P., Australian Ovarian Cancer Study Group, Mitchell, G., Fereday, S., DeFazio, A., Bowtell, D. D. L., Kristensen, G. B., Lingjærde, O. C., and Børresen-Dale, A.-L. (2013). High levels of genomic aberrations in serous ovarian cancers are associated with better survival. PLoS One, 8(1):e54356.

Bonnet, F., Guedj, M., Jones, N., Sfar, S., Brouste, V., Elarouci, N., Banneau, G., Orsetti, B., Primois, C., de Lara, C. T., Debled, M., de Mascarel, I., Theillet, C., Sévenet, N., de Reynies, A., MacGrogan, G., and Longy, M. (2012). An array CGH based genomic instability index (G2I) is predictive of clinical outcome in breast cancer and reveals a subset of tumors without lymph node involvement but with poor prognosis. BMC Med. Genomics, 5:54.

Cancer Genome Atlas Network (2012). Comprehensive molecular portraits of human breast tumours. Nature, 490(7418):61–70.

Carter, S. L., Cibulskis, K., Helman, E., McKenna, A., Shen, H., Zack, T., Laird, P. W., Onofrio, R. C., Winckler, W., Weir, B. A., Beroukhim, R., Pellman, D., Levine, D. A., Lander, E. S., Meyerson, M., and Getz, G. (2012). Absolute quantification of somatic DNA alterations in human cancer. Nat. Biotechnol., 30(5):413–421.

Cerami, E., Gao, J., Dogrusoz, U., Gross, B. E., Sumer, S. O., Aksoy, B. A., Jacobsen, A., Byrne, C. J., Heuer, M. L., Larsson, E., Antipin, Y., Reva, B., Goldberg, A. P., Sander, C., and Schultz, N. (2012). The cbio cancer genomics portal: an open platform for exploring multidimensional cancer genomics data. Cancer Discov., 2(5):401–404.

Chin, S. F., Teschendorff, A. E., Marioni, J. C., Wang, Y., Barbosa-Morais, N. L., Thorne, N. P., Costa, J. L., Pinder, S. E., van de Wiel, M. A., Green, A. R., Ellis, I. O., Porter, P. L., Tavaré, S., Brenton, J. D., Ylstra, B., and Caldas, C. (2007). High-resolution aCGH and expression profiling identifies a novel genomic subtype of ER negative breast cancer. Genome Biol., 8(10):R215.

Davison, J. M., Yee, M., Krill-Burger, J. M., Lyons-Weiler, M. A., Kelly, L. A., Sciulli, C. M., Nason, K. S., Luketich, J. D., Michalopoulos, G. K., and LaFramboise, W. A. (2014). The degree of segmental aneuploidy measured by total copy number abnormalities predicts survival and recurrence in superficial gastroesophageal adenocarcinoma. PLoS One, 9(1):e79079.

Duijf, P. H. G., Nanayakkara, D., Nones, K., Srihari, S., Kalimutho, M., and Khanna, K. K. (2019). Mechanisms of genomic instability in breast cancer. Trends Mol. Med., 25(7):595–611.

Grayson, B. L. and Aune, T. M. (2011). A comparison of genomic copy number calls by partek genomics suite, genotyping console and birdsuite algorithms to quantitative PCR. BioData Min., 4:8.

Hou, Y., Song, H., Croteau, D. L., Akbari, M., and Bohr, V. A. (2017). Genome instability in alzheimer disease. Mech. Ageing Dev., 161(Pt A):83–94.

Laddha, S. V., Ganesan, S., Chan, C. S., and White, E. (2014). Mutational landscape of the essential autophagy gene BECN1 in human cancers. Mol. Cancer Res., 12(4):485–490.

Lee, A. J. X., Endesfelder, D., Rowan, A. J., Walther, A., Birkbak, N. J., Futreal, P. A., Downward, J., Szallasi, Z., Tomlinson, I. P. M., Howell, M., Kschischo, M., and Swanton, C. (2011). Chromosomal instability confers intrinsic multidrug resistance. Cancer Res., 71(5):1858–1870.

López-Otín, C., Blasco, M. A., Partridge, L., Serrano, M., and Kroemer, G. (2013). The hallmarks of aging. Cell, 153(6):1194–1217.

McGranahan, N., Burrell, R. A., Endesfelder, D., Novelli, M. R., and Swanton, C. (2012). Cancer chromosomal instability: therapeutic and diagnostic challenges. EMBO Rep., 13(6):528–538.

Negrini, S., Gorgoulis, V. G., and Halazonetis, T. D. (2010). Genomic instability–an evolving hallmark of cancer. Nat. Rev. Mol. Cell Biol., 11(3):220–228.

Pikor, L., Thu, K., Vucic, E., and Lam, W. (2013). The detection and implication of genome instability in cancer. Cancer Metastasis Rev., 32(3-4):341–352.

Roylance, R., Endesfelder, D., Gorman, P., Burrell, R. A., Sander, J., Tomlinson, I., Hanby, A. M., Speirs, V., Richardson, A. L., Birkbak, N. J., Eklund, A. C., Downward, J., Kschischo, M., Szallasi, Z., and Swanton, C. (2011). Relationship of extreme chromosomal instability with long-term survival in a retrospective analysis of primary breast cancer. Cancer Epidemiol. Biomarkers Prev., 20(10):2183– 2194.

Song, L., Bhuvaneshwar, K., Wang, Y., Feng, Y., Shih, I.-M., Madhavan, S., and Gusev, Y. (2017). CINdex: A bioconductor package for analysis of chromosome instability in DNA copy number data. Cancer Inform., 16:1176935117746637.

Spearman, C. (1904). nthe proof and measurement of association between two things, oamerican J.

The Pan-Cancer Atlas (2022). The Pan-Cancer atlas. https://gdc.cancer.gov/about-data/publications/pancanatlas. Accessed: 2022-NA-NA.

Vargas-Rondón, N., Villegas, V. E., and Rondón-Lagos, M. (2017). The role of chromosomal instability in cancer and therapeutic responses. Cancers, 10(1).

Vijaya Sharma, S., and Batra, N. (2019). Comparative study of single linkage, complete linkage, and ward method of agglomerative clustering. In 2019 International Conference on Machine Learning, Big Data, Cloud and Parallel Computing (COMITCon), pages 568–573.

Voutsadakis, I. A. (2021). The landscape of chromosome instability in breast cancers and associations with the tumor mutation burden: An analysis of data from TCGA. Cancer Invest., 39(1):25–38.

Yurov, Y. B., Vorsanova, S. G., and Iourov, I. Y. (2019). Chromosome instability in the neurodegenerating brain. Front. Genet., 10:892.

